# EMHP: An accurate automated hole masking algorithm for single-particle cryo-EM image processing

**DOI:** 10.1101/154211

**Authors:** Zachary Berndsen, Charles Bowman, Haerin Jang, Andrew B. Ward

## 1 Introduction

The recent surge in popularity of single-particle cryo-EM as a tool for molecular structure determination alongside advances in software that have reduced the computational infrastructure needed to process single-particle datasets (Kimanius et al, 2016) have created the need for a more streamlined suite of tools to locally facilitate initial data treatment and make processing more attainable at the workstation level.

Current technical limitations inherent to the process of structure determination via single-particle cryo-EM require collecting very large data sets – often several thousands of images. This task is facilitated by automated imaging software, however downstream preprocessing steps such as quality assessment and masking of individual images are still performed manually by the researcher and can become quite cumbersome. EMHP focuses on streamlining this preprocessing stage – specifically image assessment, masking, and pick filtering in preparation for single-particle analysis.

The need for hole masking in images stems from the fundamentals of cryo-EM sample preparation and data processing. Samples are traditionally prepared for imaging by flash freezing a few microliters of solution on copper mesh grids coated in holey carbon. The hole patterns suspend particles in a thin meniscus of vitreous ice, ideal for high resolution imaging, and are often used by automated collection software to assist in exposure targeting. (Suloway et al, 2005) While images are ideally taken entirely over these holes of thin ice, it is still often necessary to collect around the edges of the holes due to low particle densities, preferential particle distribution, or contrast transfer function (CTF) estimation limitations. For these reasons, many images in a cryo-EM dataset contain sections of thick carbon support present in one or more quadrants of the image.

While automated particle selection algorithms allow for rapid selection of single particle projections from within larger images, these algorithms can have difficulty discriminating between particles on carbon and particles in ice, resulting in the inclusion of many false positives into the initial particle stack which can increase computation times and have adverse effects on downstream processing. Here we present a fast and accurate algorithm for detecting carbon supports in cryo-EM images, along with a small suite of tools for image assessment and pick filtering that allow users to preprocess their data rapidly and with minimal overhead while providing results in a format readable or easily converted to be readable by common single-particle cryo-EM processing packages. (Kimanius et al, 2016, Tang et al, 2007) Our algorithm shows improved performance over existing hole-finding algorithms and can be easily executed in a minimal or non-HPC environment making it more accessible to users.

## 2 Software Package Overview

The software included in the EMHP package is coded using Python 2.7 and relies on packages that are freely available. EMHP uses the .mrc file parser methods implemented in pyami via Appion, (Lander et al, 2009) which are included in the code repository for convenience. The package includes a Tkinter-based GUI image assessor, an implementation of the automatic hole masker and particle filter, a Tkinter-based GUI for manual hole masking, and a script that applies already computed masks to images.

### 2.1 Algorithm Description

We based our hole-finding algorithm on two main assumptions: that carbon holes are milled consistently to specifications and that edge of the holes have more detectable textural features than the vitreous ice and particles within. First, the image is decimated 10X, Gaussian filtered, and normalized via contrast stretching. (Fig. 1A) Next, a standard Sobel filter is applied, which detects the many small edges along the rim of the carbon hole. (Fig. 1B) A series of smoothing filters and thresholding operations are performed to amplify areas of concentrated signal produced during Sobel filtering, while diminishing sparse signal from particles or features in ice and on carbon. The first round of smoothing is performed on the image (Fig. 1C), and all pixels below the first threshold parameter are set to zero. (Fig. 1D) This image is then subjected to another round of smoothing and binarized using the second threshold value. (Fig 1E) We found that the first round smoothing and thresholding tends to eliminate signal from within the ice while the second round eliminates signal from within the carbon support. Finally, the image is decimated another 10X and a circle with dimensions set by the pixel size of the image and diameter of the hole is fit to the binary image with the assumption that the largest match will be along the carbon edge. The fitting is performed exhaustively Θ(n^2^), but is sped up considerably by the decimation steps. The coordinates of the best fit circle are then extrapolated back to the original image and the mask is shifted toward or away from the edge of the hole based on user input. This final binary mask is used to filter out particle picks that lie on top of the carbon support (Fig. 1F). We have also included an option to filter picks that are within a user defined distance from the edge of the image, usually set to the desired box size used during single particle processing. Users can adjust several filter and threshold parameters to tune performance for their data set, and we have found that the most impactful parameter is the intensity cutoff used in the second thresholding step, which controls the width and shape of the final binary edge used for circle fitting. Descriptions of all parameters and examples of their effects can be found in supplementary materials.

**Fig. 1.**
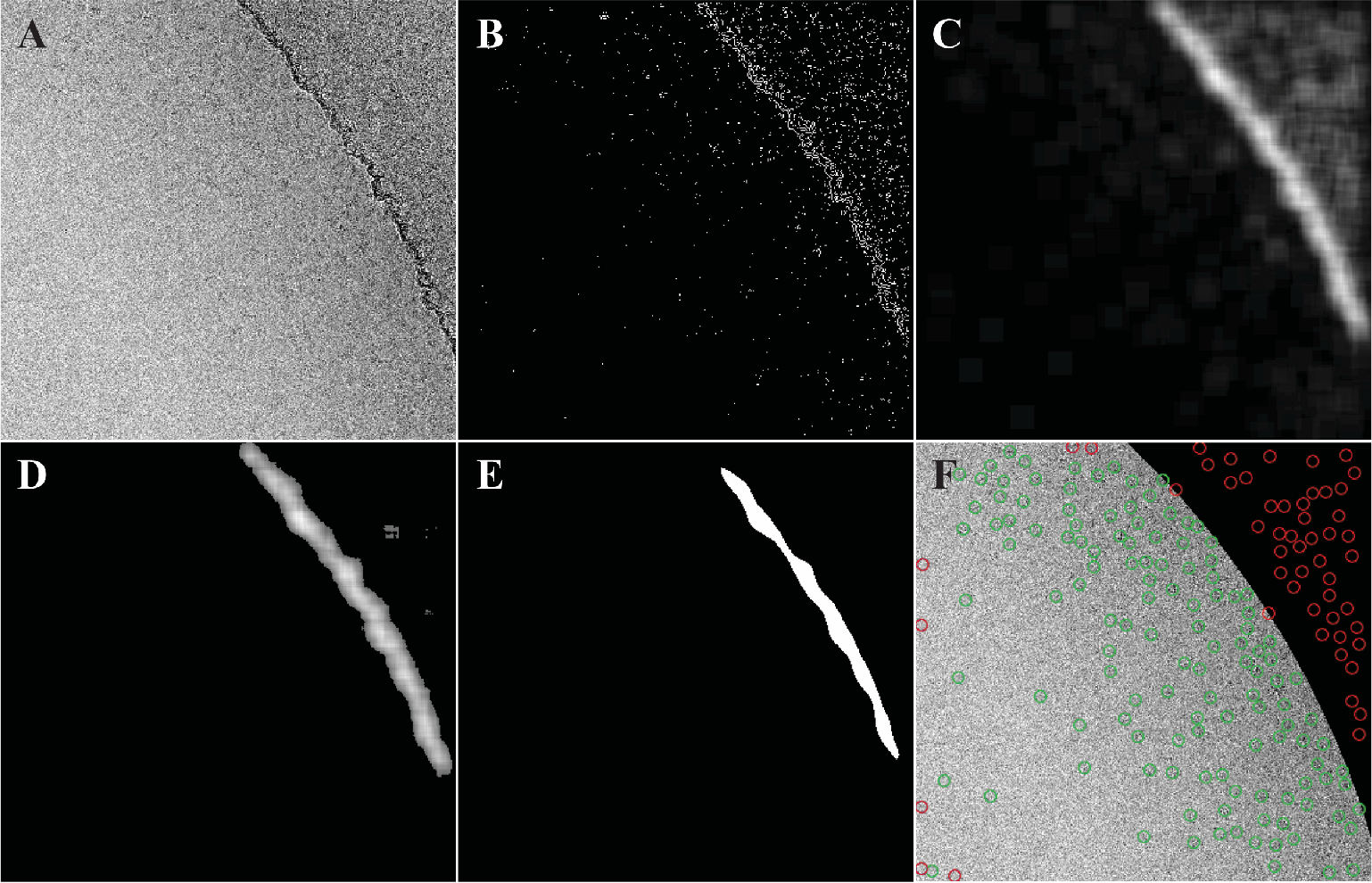
EMHP automasking example. The image after (A) Gaussian filter and contrast stretching, (B) after applying a Sobel filter, (C) after one round of edge amplification by radial summing, (D) after applying a user-defined threshold and (E) after another round of edge amplification and binary thresholding. (F) The final image after circle fitting, masking, and pick filtering. Green single particle picks are included, while red picks are excluded due to the mask or edge proximity.

### 2.2 Test Dataset and Methods

For a concrete evaluation of the algorithm’s effectiveness, we conducted two performance tests on a set of 100 images chosen at random from a previously published dataset covering a defocus range of -1.4 to -3.6µm (Lee et al, 2016). We compare performance of the most commonly used automasker, em_hole_finder, currently incorporated into the Appion web-based software suite (Lander et al, 2009) with the EMHP automasking algorithm. To do this, masks were carefully constructed manually and a set of particle picks generated with DoG Picker (Voss et al, 2009) were assigned to either ice or carbon. Next, both algorithms were used to classify the same set of picks and the sensitivity (true positive rate), specificity (true negative rate), and average runtime per image were calculated for each (Table 1). These 100 test images have been included in the code repository for testing and benchmarking.

**Table 1.**
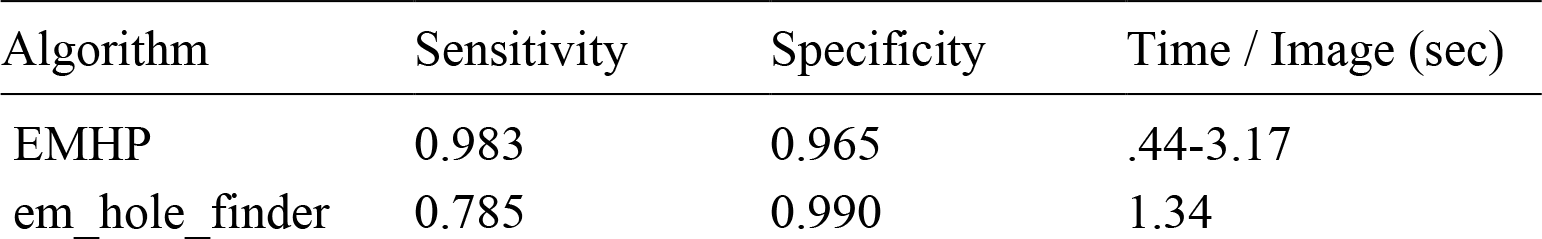
Benchmark comparisons using em_hole_finder and EMHP. EMHP per-image calculation times reported as a range with the mini-mum bound representing the program run in multithreaded (16x) mode. This mode is not available in em_hole_finder at time of submission.

To further justify the utility of masking we performed a single round of 2D-classification using GPU-accelerated RELION-2.0 on the full particle stack from the test dataset, this same stack filtered using manually produced masks, and a stack that is the same size as the masked stack made from a random subset of particles. Around 40% of the particle picks were filtered out during masking, and it is therefore not surprising that 2D-classification ran 1.8X faster, however the masked data set still ran 1.1X faster than the unmasked dataset of equivalent size, illustrating that the inclusion of false positives does slow down 2D-classification. Additionally, 36% of the particles selected after 2D-classification from the unmasked data set were false positive picks on carbon, implying that 2D-classification alone cannot completely sort out picks on carbon.

## 3 Conclusions

The EMHP automasking algorithm shows improved sensitivity com-pared to em_hole_finder. This highlights the main shortcoming of em_hole_finder, the tendency to over-mask thereby excluding a sub-stantial number of true positives from the initial particle stack. The two algorithms performed equivalently in terms of specificity.

We would like to note that over the course of testing and extensive in-house use, we found that the EMHP algorithm has trouble with edge detection in two specific situations: when carbon edges are poorly manufactured (overly jagged or un-circular), or when particle density at the very edge of the carbon holes is excessively high. In comparison, we observe that em_hole_finder does not have additional issues with poorly manufactured edges, but tends to over-mask when presented with dense patches of particles along the edges. To get around these performance limitations, the EMHP suite also includes tools for sorting images before masking and applying manually placed circular masks.

Rapid advances in single-particle cryo-EM data processing software and hardware capabilities are bringing state-of-the-art structure deter-mination capabilities to the desktop. EMHP provides a simple python package for single-particle cryo-EM image preprocessing that is ac-cessible to users of all experience levels while requiring minimal com-putational overhead.

